# Intranasal Insulin Mediates Neurodegeneration in Diabetic Retinopathy via Regulation of Inflammatory and Apoptotic Pathways

**DOI:** 10.1101/2025.07.23.666398

**Authors:** Sally S. Ong, Joanne Konstantopoulos, Mallory K. Suarez, Joseph Rigdon, Jian-Xing Ma, Rebecca M. Sappington

**Author notes:** Corresponding Author: Sally S. Ong, MD Assistant Professor of Ophthalmology Wake Forest University School of Medicine 1 Medical Center Blvd, Winston-Salem, NC, 27157.

## Abstract

Neurodegenerative changes predominate in early stages of diabetic retinopathy but effective therapies are lacking. Insulin treatment decreases neurodegeneration and intranasal insulin has been shown to reach the central nervous system in neurodegenerative diseases like dementia. We tested the hypothesis that intranasal insulin can decrease retinal neurodegeneration using the C57BL/KsJ-db/db transgenic diabetic (*db/db*) mouse model. Compared to the non-diabetic wildtype mice given intranasal saline, we observed decreased electroretinogram b-wave and oscillatory potential amplitudes in *db/db* mice treated with intranasal saline but not in the db/db mice treated with 2 units of intranasal insulin daily over 10 weeks. When compared to the non-diabetic intranasal saline control, we also observed decreased outer retinal thickness in the *db/db* mice given intranasal saline but this effect was attenuated in the *db/db* mice treated with intranasal insulin. GFAP immunoreactivity and caspase cell count were similarly elevated in the *db/db* mice treated with intranasal saline but not intranasal insulin. Mean blood glucose measurements increased 30 minutes after both intranasal saline and insulin treatment. Transcriptomic analysis revealed downregulation of inflammatory and apoptotic genes in the retina of *db/db* mice treated with intranasal insulin when compared to saline. In summary, treatment with intranasal insulin prevents the depression of b-waves and oscillatory potentials, decreases the attenuation of outer retinal thickness, reduces caspase cell count and GFAP immunostaining, and downregulates the transcription of inflammatory and apoptotic genes in the retina of *db/db* mice without exerting peripheral glucose lowering effects. Taken together, our results suggest that intranasal insulin can reduce neurodegeneration in diabetic retinopathy by improving retinal neuronal function, decreasing reactive gliosis and cell death, and modulating the expression of inflammatory and apoptotic genes.

## Introduction

Diabetic retinopathy (DR) is the most common microvascular complication in diabetes and the most frequent cause of acquired blindness in working age adults worldwide.^1^ In 2020, DR was estimated to affect over 100 million adults globally and of these, almost 29 million had vision-threatening complications.^1^ In recent years, diabetic neurodegeneration has been shown to be an important component of DR that may occur independently or interdependently with vascular changes^2^ and no therapy exists yet for diabetic neurodegeneration. DR is increasingly recognized as a disorder of the neurovascular unit, which refers to the interdependency of neurons, glia and vasculature to maintain normal retina function.^3^ Pathologic changes of the neurovascular unit in DR can affect both the vasculature and the neural retina, and these changes can occur via activation of apoptosis and inflammatory pathways.^4^

Insulin is a potent anabolic hormone that has been found to rescue these neurons and glia from apoptotic cell death.^5–7^ Systemic, intravitreal and subconjunctival administration of insulin has been shown to restore prosurvival insulin receptor and Akt kinase activity and to decrease apoptotic cell death associated with diabetes.^6,8,9^ Insulin has also been shown to prevent the activation of inflammatory pathways by suppressing major pro-inflammatory transcription factors like NF-κβ, activator protein-1 (AP-1) and early growth response-1 (EGR-1).^10,11^ However, there are numerous challenges associated with current routes of insulin delivery. Insulin has a short half-life in the plasma,^12^ and it is difficult to administer sufficient systemic insulin to reduce the risk of retinopathy without causing hypoglycemia.^13^ Topical eye drops are not effective for treatment of retinal diseases due to corneal and conjunctival barriers, and rapid precorneal tear loss.^14,15^ Intravitreal and subconjunctival injections are invasive and carry risks of infection.^9,16^ In contrast, intranasal insulin administration is non-invasive, can be easily self-administered, avoids hepatic first-pass elimination and has been shown to reach the central nervous system within minutes without raising peripheral insulin or causing hypoglycemia.^17,18^

Intranasal insulin has been extensively studied in Alzheimer’s disease and has been shown to improve cognitive performance.^19^ Insulin resistance is thought to contribute to cognitive decline, and low insulin levels in the central nervous system may be caused by impaired insulin transport through the blood brain barrier. In contrast, insulin delivered intranasally has been shown to cross the blood brain barrier and was detected in the brainstem, cerebellum, substantia nigra/ventral tegmental area, olfactory bulb, striatum, hippocampus and thalamus/hypothalamus.^20^ We hypothesized that intranasal insulin can similarly cross the blood retinal barrier leading to increased insulin levels in the retina. We therefore explored the effects of intranasal insulin delivery on retinal neuronal function, apoptosis and inflammation in a mouse model of diabetes.

## Methods

### Animals

Two-month-old C57Bl/6J (wild-type, WT) and B6.BKS(D)-Lepr^db^/J (*db/db*) male mice (27-29g, 44-50g, resp.) were purchased from The Jackson Laboratory (Bar Harbor, ME, USA). Animals were housed under standard conditions with *ad libitum* food and water and maintained on a 12-h light/dark cycle. Mice were euthanized by cervical dislocation under Isoflurane vapor (5%) anesthesia. The right eye of each animal was rapidly removed and flash frozen on dry ice and then stored at -80°C. The left eye was drop fixed into 4% paraformaldehyde/1X phosphate buffered saline (PBS) (both from Fisher, Scientific, Atlanta, GA) and stored at 4°C. All studies were carried out following the Guide for the Care and Use of Laboratory Animals, 8^th^ edition and approved by the Wake Forest School University School of Medicine Animal Care and Use Committee.

Before treatments began, all mice underwent a series of baseline measurements, including weight (repeated periodically to monitor animal health and adjust anesthesia for ERG), blood-glucose (BG) levels (Contour Next EZ Blood Glucose Monitoring System and test strips), and dark-adapted electroretinogram (ERG) and oscillatory potentials (OP) (Celeris by Diagnosys, LLC).

### Intranasal Insulin Administration

All WT mice received intranasal administration of 0.9% bacteriostatic saline, while the DB/DB mice were split into two groups, where an equal number received saline or 2 units (Humulin R, U-100, Eli Lilly) of intranasal insulin. All animals received a total of 20ul (10ul per nostril) of liquid, daily, for 10 weeks. For intranasal administration, mice were scruffed in a vertical position while liquid was slowly expelled into each nostril (10-20sec) using a 10ul Denville pipet. Mice were held in this position for at least 30 seconds after administration (with normal breathing) so that liquid could not be wiped away during grooming.

### Blood Glucose (BG) Levels

Within treatment groups, mice were divided into two groups such that BG levels were taken every other week per group throughout the study, for a total of 5, in-study measurements per mouse. For the measurement, food was removed from cage and BG reading was taken. 30 minutes later, BG levels were taken again, and food was replaced. BG was obtained by briefly warming tail on a Deltaphase® isothermal pad, after which a sterile 26Ga needle was inserted into the tail vein to collect approximately 10-20ul of blood and placed on test strip.

### Dark-Adapted electroretinogram (ERG) and oscillatory potentials (OP)

For dark-adapted ERG and OP testing, the dark cycle was extended so that mice were housed in the dark for at least 16 hours before running the measurement. The mice were transported to test room under a blackout curtain covered with black plastic and allowed to acclimate to testing room where a red light was used to run experiment. Mice were anesthetized with Ketamine (80-100mg/kg) and Xylazine (5-10mg/kg) (both from Patterson Veterinary Supply, Charlotte, NC). 1% Tropicamide (Somerset Therapeutics LLC) was used to dilate pupils and 0.5% Proparacaine Hydrochloride (Bausch and Lomb) added for numbing. After mice were unresponsive to toe pinch, they were placed on the heated (37°C) stage of ERG system and Systane lubricating gel (Alcon [Hypomellose 0.3%]) placed onto eyecups. These were placed such that gel made contact between eyes and eye cups, without exerting force into eyes. A series of flashes of white light at an intensity of 1.0cd sec/m^2^ at a flash frequency of 1Hz with an inner sweep delay of 500msec were given to each eye. After the experiment, GenTeal® (Alcon [Dextran70 0.1%, Glycerin 0.2%, Hypomellose 0.3%]) drops were placed on eyes, animals were monitored until they fully recovered from anesthesia and placed back in housing. ERG testing was repeated at the end of the study.

### Tissue Preparation, Immunohistochemistry and Microscopy

The left eyes were cryoprotected in a series of sucrose (Fisher Scientific, Atlanta, GA) dilutions (10%, 20%, 30%) in 1XPBS with 0.2% sodium azide (Sigma Aldrich, Saint Louis, MO), kept at 4°C until tissue sunk at each concentration and cut on a Leica cryostat at 10um in the naso-temporal plane. Slides were kept at -80°C until immunohistochemistry (IHC) was performed. Briefly, tissue was removed from - 80°C freezer and dried at room temperature for at least a half hour, put into auto-quench solution (1% sodium borohydride in 1X PBS (Fisher Scientific, Atlanta, GA) and washed 3 times at 5 min each with 1X PBS, while shaking for 30 min. Following the blocking step (5% normal donkey serum (NDS) (Fisher Scientific, Atlanta, GA), 0.1% Triton-X (Sigma Aldrich, Saint Louis, MO) and 1X PBS for 2 hrs at RT, shaking, primary antibody cocktails were made at respective concentrations in 3% NDS, 0.1% Triton-X and 1XPBS/0.02% sodium azide, applied to tissue and left at 4^0^C overnight. On day two, primary antibody was removed in a series of 4 washes in 1X PBS at RT, shaking. Secondary antibody cocktails were made with 1% NDS, 0.1% Triton-X and 1XPBS, and incubated for 2 hrs at RT, shaking. After 4, 10 min washes in PBS, shaking at RT and one wash in ultrapure water for 10 min at RT, shaking, tissue was coverslippped with Fluoromount-G with DAPI (Fisher, Scientific, Atlanta, GA) and dried overnight at RT. On day three, slides were sealed with Cytoseal 60 (Fisher, Scientific, Atlanta, GA) and stored for long-term use at 4^0^C.

Immunostained slides were then imaged using confocal microscopy (C2 Nikon Ti2) at 20X. Comparative images were taken using identical laser power and gain settings. All tissue samples were imaged at 5 different locations on the retina (P1=nasal end, C1=between P1 and optic nerve, C2=approximate location of optic nerve, C3, between optic nerve and P2 and P2=temporal end). Only images from C1, C2 and C3 were analyzed. Measurements were taken within the neural retina as defined from the internal limiting membrane to the outer border of the outer nuclear layer. For cell counting, the cell counter feature on ImageJ was utilized. For intensity, the integrated intensity measurement feature on ImageJ was used.

**Table 1.** List of antibodies used

| Antibody | Epitope | Host | Concentration | Part Number | Vendor |
| --- | --- | --- | --- | --- | --- |
| Glial Fibrillary Acidic Protein | -- | Rb | 1:500 | Z033401-2 | Agilent Technologies |
| Cleaved Caspase-3 | Asp175 | Rb | 1:400 | 9661 | Cell Signaling |
| Recombinant Anti-Insulin | EPR17359 | Rb | 1:1000 | ab181547 | abcam |
| Click-iT™ Plus TUNEL assay | -- | -- | -- | C10619 | Invitrogen |

### Retinal layer thickness measurements

For measured layer thickness, one slide per animal was stained with Toluidine blue O (Fisher, Scientific, Atlanta, GA) and imaged using light microscopy. Using ImageJ, inner retinal thickness was measured from the internal limiting membrane to the outer border of the inner nuclear layer. The outer retinal thickness was measured from the outer border of the inner nuclear layer to the outer border of the outer nuclear layer.

### RNA Sequencing

Total RNA was isolated from frozen retina tissues of db/db mice treated with saline and 2U insulin. The RNA-Seq was performed at the WFUSM Cancer Genomics Shared Resource at. Analysis of RNA-seq data was performed at the WFUSM Bioinformatics Shared Resource. Sequencing files were trimmed and cleaned with Trimmomatic v0.33 and mapped with STAR v2.5.3 before being collected into reads per gene by featureCounts v1.4.5. EdgeR v4.2.0 was used for differential expression analysis. Gene set enrichment analysis was performed with cluster Profiler v4.16.0 with gene sets provided by msigdbr v25.1.0. Gene set enrichment was performed using Hallmark gene sets and an adjusted p-value cutoff of 0.05 after applying the Benjamini-Hochberg false discovery rate adjustment. A network graph was produced using two genes making up the leading edges of two gene sets found to be significantly inhibited in our analysis.

### Statistical analysis

Generalized linear mixed effects models were used to conduct within (over time, where appropriate) and between-group comparisons in outcomes. In these models, fixed effects were included for group (and time, and time by group interaction, where appropriate) and random effects were included for animal (and nested eye within animal, where appropriate). Prior to applying the mixed models, outliers were removed. Following standard statistical practice, an observation was deemed an outlier when it was either one-and-a-half times the interquartile range below the first quartile or one-and-a-half times the interquartile range above the third quartile. In hypothesis testing using these models, Kenward-Roger denominator degrees of freedom adjustments were made. Estimated marginal means and confidence intervals from these models were displayed using bar plots to visualize the findings. Statistical significance was set at p<0.05. Statistical analyses were performed using R version 4.3.2.

## Results

The B6.BKS(D)-*Lepr^db^*/J mice with type 2 diabetes used throughout this study are subsequently referred to as *db/db* mice. All saline and insulin treatments mentioned herein were delivered intranasally. To verify that intranasal insulin did not confer its effects by lowering systemic blood glucose, we measured peripheral blood glucose levels before and 30 minutes after administering intranasal treatment (Figure 1). While peripheral glycemic levels did not differ before and after treatment for the C57BL/6 saline mice (p=0.6088, n=9 mice), they increased post treatment for both the *db/db* saline [34.04 (95% CI: 4.22, 63.86); p=0.0254, n= 10 mice] and *b/db* insulin [41.93 (95% CI: 11.81, 72.05); p=0.0065; n = 10 mice] groups, therefore demonstrating that intranasal insulin did not act peripherally to reduce blood glucose levels.

**Figure 1.**
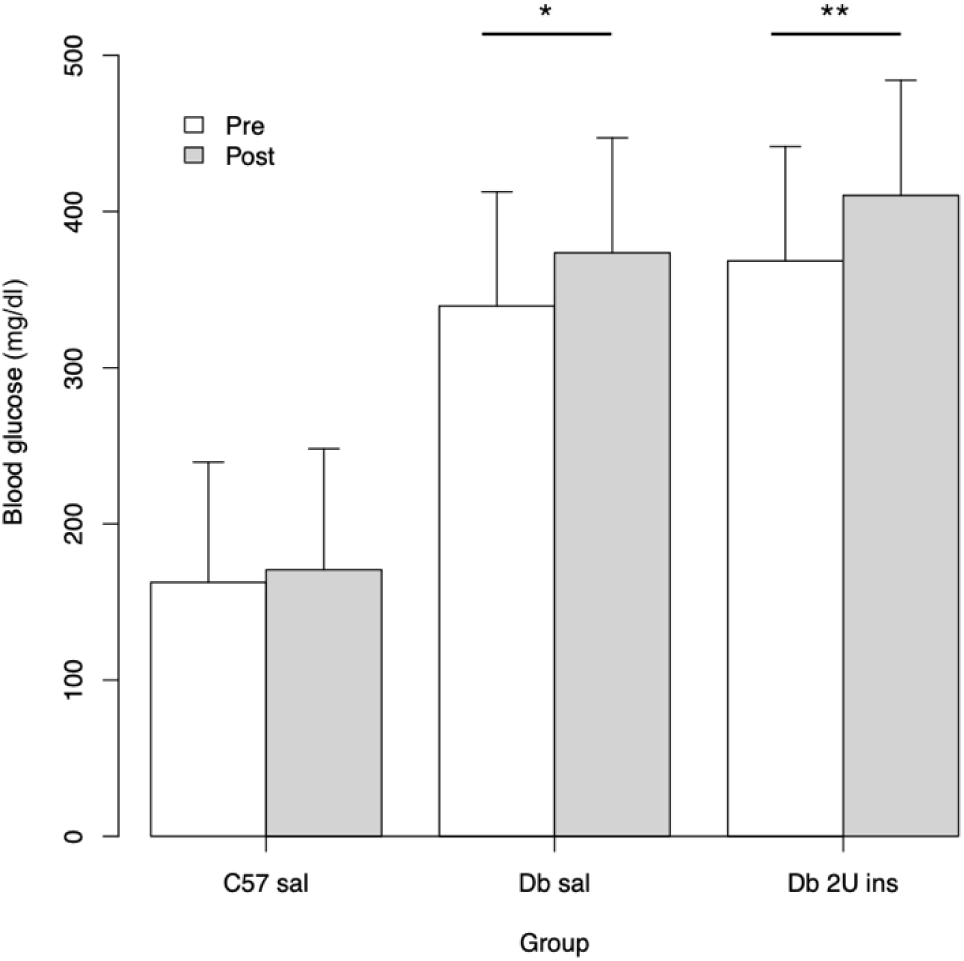
Peripheral blood glucose measurements before and after intranasal treatment. Mean glucose measurements did not differ before or after intranasal treatment in the non-diabetic C57 group (n=9; p=0.6) but increased post treatment for the diabetic *db/db* mice treated with either intranasal saline (n=10; p=0.03) or insulin (n=10; p=0.007). *P<0.05; **P<0.01, by generalized linear mixed effects models. Data represent the estimated marginal means +/- CI.

Using scotopic ERG testing, we compared changes in a-wave and b-wave amplitudes before and after 10 weeks of intranasal insulin (*db/db* insulin, n=4 eyes) versus saline therapy (C57 saline, n=4 eyes; *db/db* saline, n=4 eyes) (Figure 2). Except for a reduction in a-wave amplitude after treatment in the *db/db* insulin group at a light intensity of 0.01 cd.s/m2 [1.72 (95% CI: 0.44, 3.01); p=0.0107], there was no difference in a-wave amplitudes pre and post treatment for the three experimental groups at 0.01 cd.s/m2 (C57 saline: p=0.8363; *db/db* saline: p=0.4568), 0.1 cd.s/m2 (C57 saline: p=0.7305; *db/db* saline: p=0.8922; *db/db* insulin: p=0.6120), or 1.0 cd.s/m2 (C57 saline: p=0.5282; *db/db* saline: p=0.2175; *db/db* insulin: p=0.4967).

**Figure 2.**
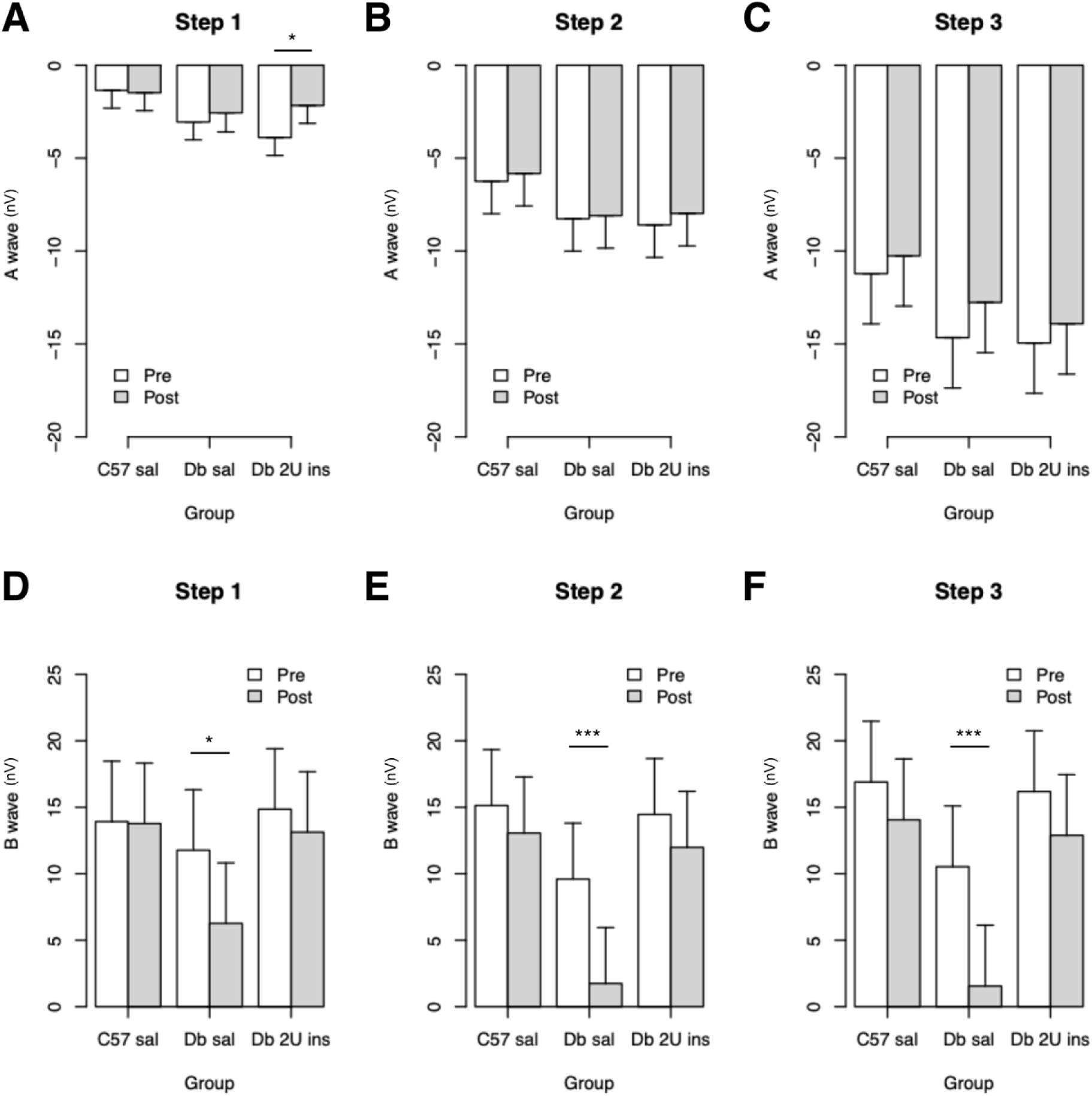
ERG a and b-waves at 0.01 cd.s/m2 (Step 1), 0.1 cd.s/m2 (Step 2) and 1.0 cd.s/m^2^ (Step 3) before and after intranasal treatment. A. A-wave amplitude improved pre and post treatment in the diabetic insulin group (p=0.0107, n=4) at Step 1 light intensity. Otherwise, there was no difference before and after treatment in the non-diabetic (n=4) and diabetic (n=4) groups treated with saline. B. No difference in a-wave amplitude pre and post treatment was noted in any of the three groups studied at Step 2 light intensity. C. There was no difference between any of the groups studied in the a-wave amplitude pre and post treatment. D. The diabetic group treated with saline (n=4) had decreased b-wave amplitudes post treatment (p=0.0276), but this decrease was not seen in the diabetic group treated with insulin (n=4) or the non-diabetic group treated with saline (n=4) at Step 1 light intensity. E. B-wave amplitudes decreased in the diabetic saline group after treatment (p=0.0005) but not in the diabetic insulin or non-diabetic saline groups at Step 2 light intensity. F. Similarly, B-wave amplitudes were reduced after treatment in the diabetic saline group (p=0.0005) but no change post treatment was observed in the diabetic insulin or non-diabetic saline groups at Step 3 light intensity. *P<0.05; **P<0.01,**P<0.001 by generalized linear mixed effects models. Data represent the estimated marginal means +/- CI.

While there was no change in b-wave amplitudes pre and post treatment in the C57BL/6 saline group at 0.01 cd.s/m2 (p=0.9509), 0.1 cd.s/m2 (p=0.3007) or 1.0 cd.s/m2 (p=0.2145), there was a consistent reduction in b-wave amplitudes in the *db/db* saline groups across all three light intensities at 0.01 cd.s/m2 [-5.51 (95% CI:-10.35, -0.66); p=0.0276], 0.1 cd.s/m2 [- 7.86 (95% CI: -11.89, -3.83); p=0.0005] and 1.0 cd.s/m2 [-8.97 (95% CI:-13.57, -4.38); p=0.0005]. In comparison, in the *db/db* group treated with insulin, no difference in b-wave amplitudes was observed at 0.01 cd.s/m2 (p=0.4692), 0.1 cd.s/m2 (p=0.2177) or 1.0 cd.s/m2 (p=0.1522). Oscillatory potential amplitudes also did not change pre and post treatment for the C57BL/6 saline group at 0.01 cd.s/m2 (p=0.2867), 0.1 cd.s/m2 (p=0.1909) or 1.0 cd.s/m2 (p=0.2779), or for the *db/db* saline group at 0.01 cd.s/m2 (p=0.0871), but decreased after treatment in the *db/db* saline group at 0.1 cd.s/m2 [-72587.47 (95% CI: -128346.48, -16828.47); p=0.0129] and 1.0 cd.s/m2 [-88817.02 (95% CI: -161329.89, -16304.14); p=0.0185] (Figure 3). For *db/db* mice treated with insulin, oscillatory potential amplitudes did not change after treatment at 0.01 cd.s/m2 (p=0.3802), 0.1 cd.s/m2 (p=0.5252) or 1.0 cd.s/m2 (p=0.4270).

**Figure 3.**
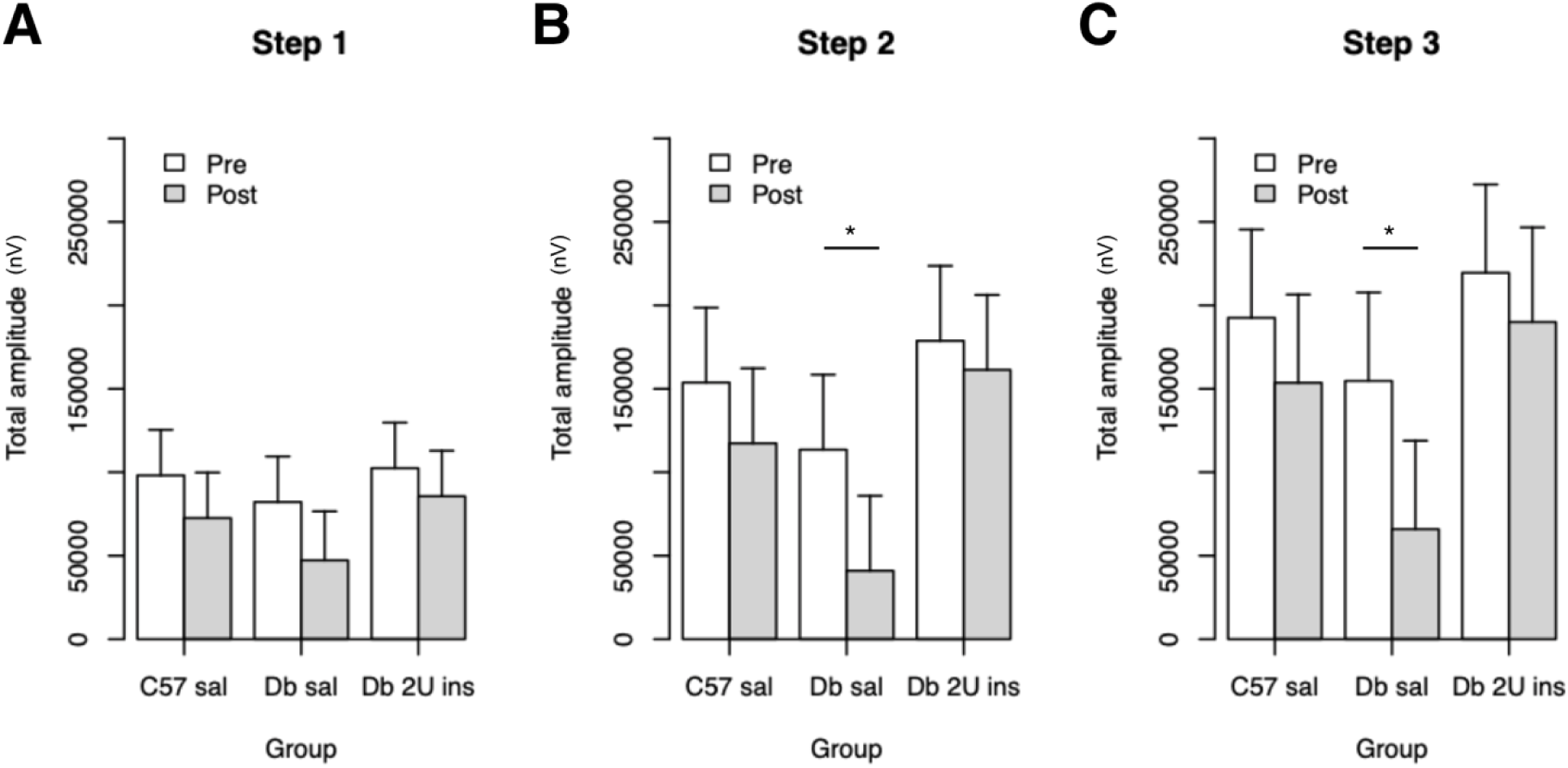
Oscillatory potentials (OP) at 0.01 cd.s/m2 (Step 1), 0.1 cd.s/m2 (Step 2) and 1.0 cd.s/m^2^ (Step 3) light intensity before and after intranasal treatment. A. No difference pre and post treatment was found in any of the three groups studied at Step 1 light intensity. B. At Step 2 light intensity, OP amplitudes decreased after treatment in the diabetic saline group (p=0.0129) but not in the non-diabetic saline or diabetic insulin group. C. At Step 3 light intensity, there was a reduction in OP amplitudes post treatment in the diabetic saline group (p=0.0185) but not in the non-diabetic saline or diabetic insulin group. *P<0.05 by generalized linear mixed effects models. Data represent the estimated marginal means +/- CI

We then examined retinal thickness across groups and found that compared to the C57BL/6 saline mice (n=9 eyes), the inner retinal thickness was reduced in both the *db/db* saline [-20.3 (95% CI:-32.14, -8.46); p=0.0016; n= 8 eyes] and *db/db* insulin -17.21 (95% CI: - 28.69, -5.73); p=0.0048; n= 9 eyes] groups (Figure 4). There was no difference in inner retinal thickness between the *db/db* saline and *db/db* insulin groups (p=0.5898). In comparison, when compared to C57BL/6 saline, outer retinal thickness was reduced in *db/db* saline -11.66 (95% CI: -21.12, -2.2); p=0.0175] but not in the *db/db* insulin (p=0.0944) group. No difference in outer retinal thickness was observed between the *db/db* saline and *db/db* insulin groups (p=0.3996).

**Figure 4.**
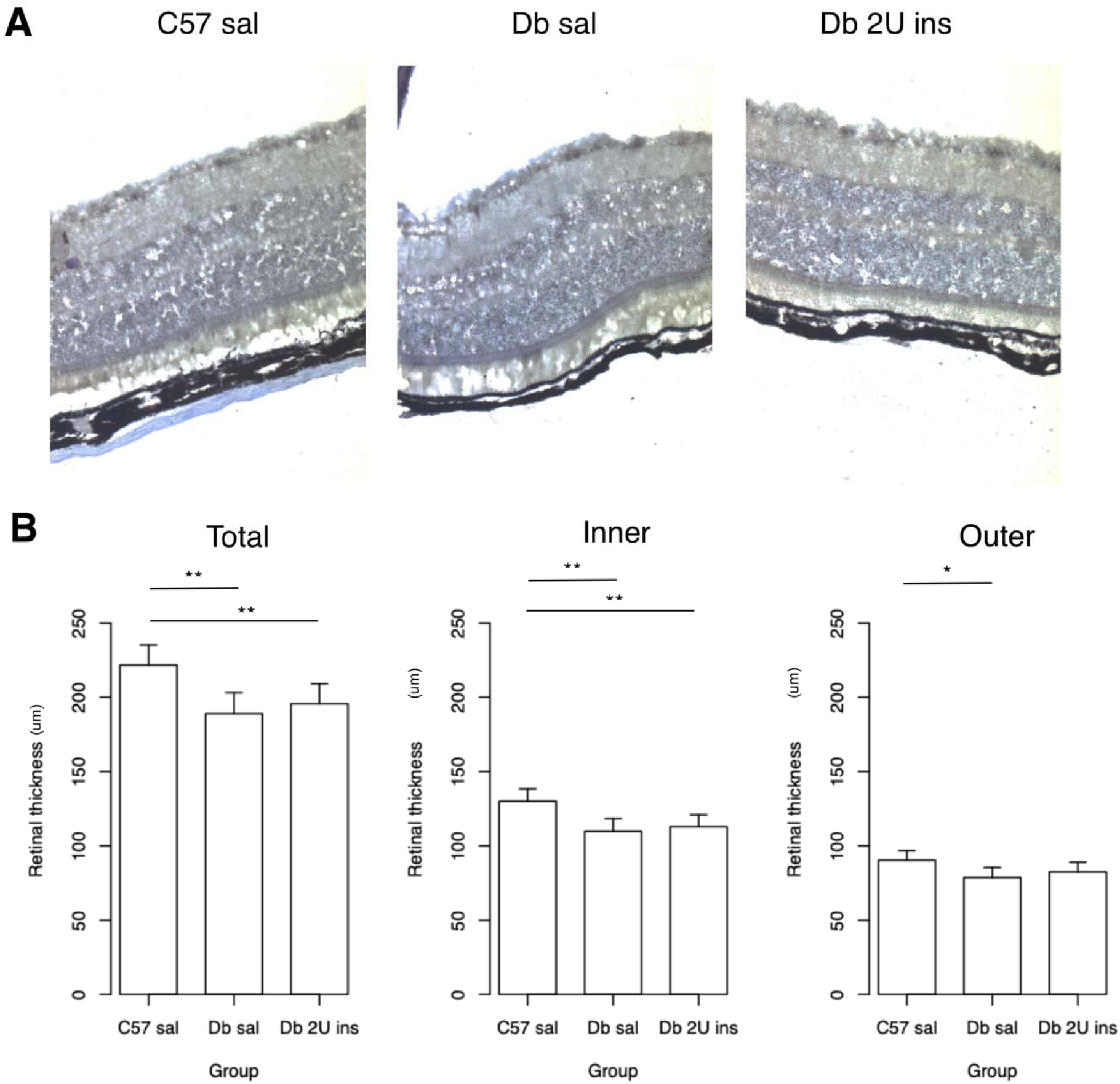
Retinal thickness measurements using retinal sections stained with Toluidine blue O. A. Total retinal thickness was decreased in both the diabetic saline (p=0.0019) and diabetic insulin groups (p=0.0092) compared to the non-diabetic control. B. The inner retinal thickness was also reduced in the diabetic saline (p=0.0016) and diabetic insulin (p=0.0048) groups compared to the non-diabetic saline group. C. Outer retinal thickness was only decreased in the diabetic saline (p=0.0175) group compared to the non-diabetic saline group, but not in the diabetic insulin group. *P<0.05 **P<0.01 by generalized linear mixed effects models. Data represent the estimated marginal means +/- CI

The pattern of b-wave and oscillatory potential amplitude reductions in the *db/db* group treated with saline but not in the *db/db* group treated with insulin was further explored by comparing GFAP expression in the retina. Representative images showing GFAP expression in each group are shown in Figure 5. Compared to C57BL/6 mice treated with saline [n=9 eyes], *db/db* mice treated with saline had an increased GFAP intensity density [0.35 (95% CI: 0.10, 0.61); p=0.0082; n= 8 eyes] while the *db/db* mice treated with insulin had reduced GFAP intensity density [-0.36 (95% CI: -0.6, -0.11); p=0.0063; n= 9 eyes]. Compared to the *db/db* saline eyes, GFAP intensity density was also decreased in the *db/db* insulin eyes [-0.71 (95% CI: -0.96, -0.46); p<.0001].

**Figure 5.**
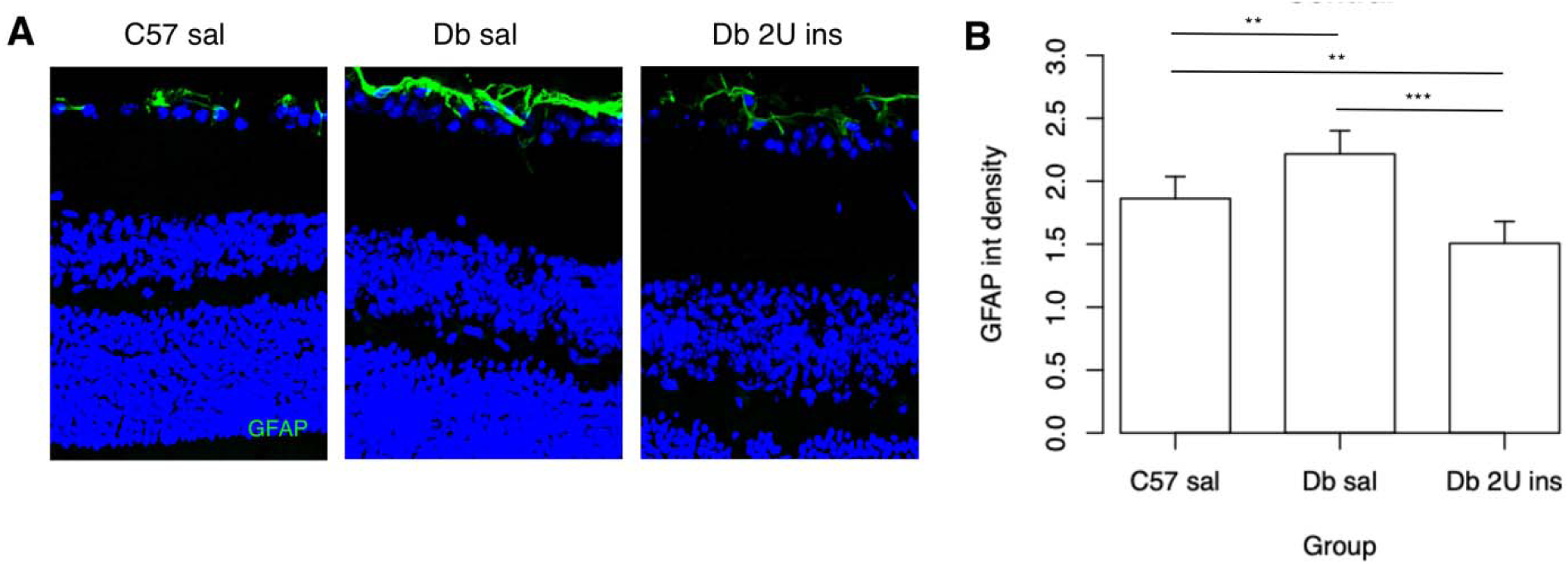
A. Glial fibrillary acidic protein (GFAP) immunostaining was measured as intensity density and compared between groups. B. GFAP staining was higher in the diabetic saline group compared to both the non-diabetic saline (p=0.0082) and diabetic insulin (p<0.0001) groups. GFAP immunoreactivity was also decreased in the diabetic insulin group when compared to the non-diabetic saline group (p=0.0063). **P<0.01 ***P<0.001 by generalized linear mixed effects models. Data represent the estimated marginal means +/- CI

Next, caspase cell count was compared between the three experimental groups. Representative images of caspase 3 expression are shown in Figure 6. Compared to the C57BL/6 saline group (n=8 eyes), caspase cell count was increased in the *db/db* saline group [3.89 (95% CI: 0.59, 7.18); p=0.0227; n= 7 eyes] but did not differ in the *db/db* insulin group (p=0.3384; n=9 eyes). There was no difference in caspase cell count between the *db/db* mice treated with saline or insulin (p=0.1373). TUNEL cell count was also examined in parallel (images not shown). There was a similar trend of TUNEL cell count being higher in the *db/db* saline (n=8 eyes) compared to the C57BL/6 saline (n=9 eyes) group although this did not reach statistical significance (p=0.0510). There was no difference in TUNEL cell count between the C57BL/6 saline and *db/db* insulin (n=8 eyes) groups (p=0.2334) or between the *db/db* saline and *db/db* insulin groups (p=0.4031).

**Figure 6.**
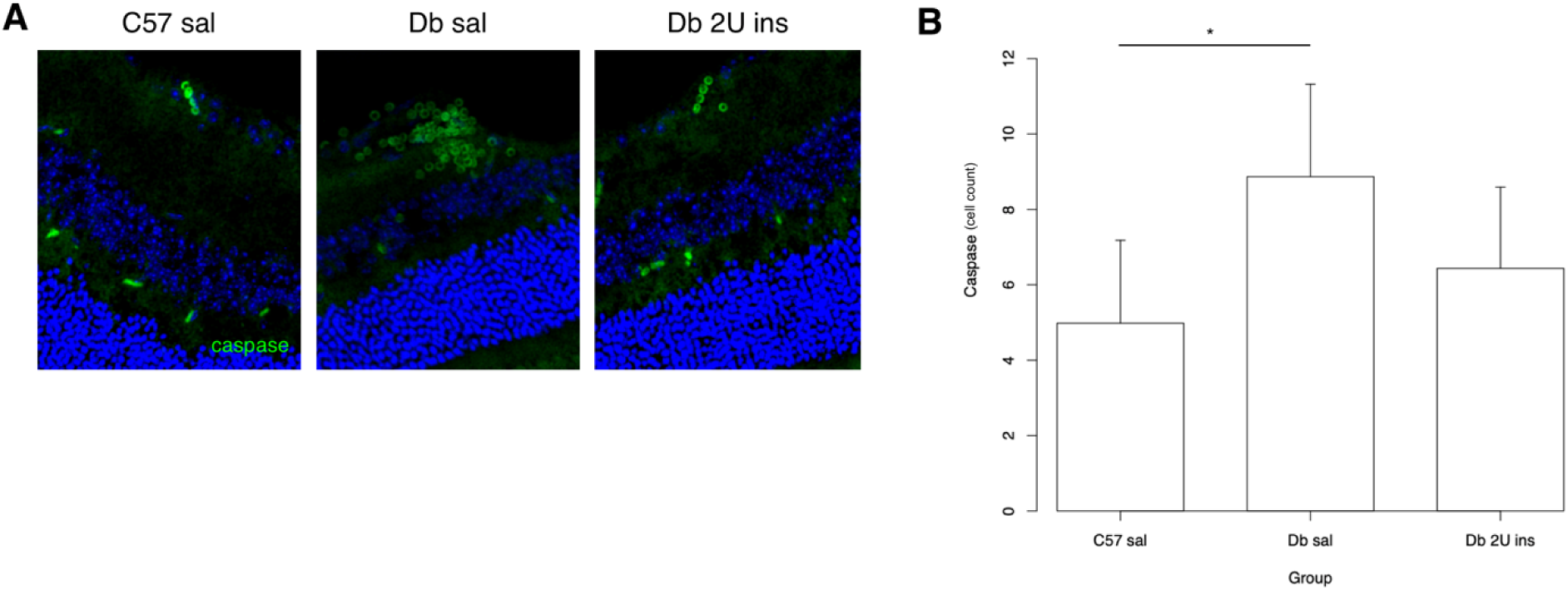
A. Cleaved caspase-3 cell count was measured and compared between groups. A. The caspase cell count in the diabetic saline group was higher than that for the non-diabetic saline group (p=0.0227). There was no difference between the diabetic group treated with intranasal insulin compared to the non-diabetic group treated with intranasal saline (p=0.3384). *P<0.05 by generalized linear mixed effects models. Data represent the estimated marginal means +/- CI

Next, we measured insulin immunofluorescent staining in the retinas of the three experimental groups to verify delivery of intranasal insulin to the retina (Figure 7). While there was no difference in insulin intensity density between the two groups treated with saline (p=0.3978), there was an increase in insulin immunofluorescence in the *db/db* insulin (n=9 eyes) group compared to both the C57BL/6 saline [8.09 (95% CI: 5.09, 11.09); p<.0001; n= 9 eyes] and *db/db* saline [6.85 (95% CI: 3.72, 9.98); p=0.0002; n= 8 eyes] groups.

**Figure 7.**
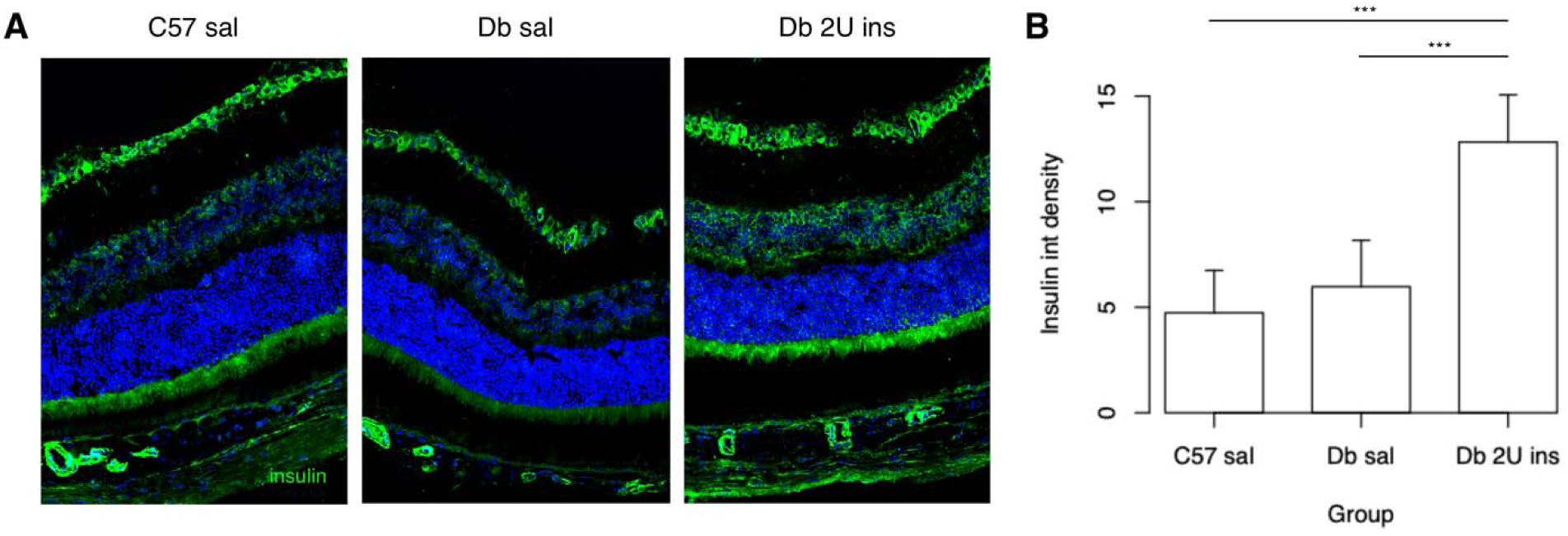
Insulin immunoreactivity was measured by intensity density from the internal limiting membrane to the external limiting membrane. B. There was a higher insulin intensity density in the diabetic insulin group compared to both the non-diabetic saline (p<0.0001) and diabetic saline (p=0.0002) groups. ***P<0.001 by generalized linear mixed effects models. Data represent the estimated marginal means +/- CI

Lastly, we used gene set enrichment analysis to determine differences in gene expression between the *db/db* mice treated with saline (n=5 eyes) versus insulin (n=5 eyes) (Figure 8). When compared to *db/db* mice treated with saline, gene set enrichment showed significant downregulation of gene sets involved in these ten pathways in the *db/db* mice treated with insulin - estrogen response late, P53 pathway, IL2 stat5 signaling, estrogen response early, interferon alpha response, allograft rejection, epithelial mesenchymal transition, xenobiotic metabolism, G2M checkpoint and apoptosis. The network graph showed that treatment with insulin downregulated expression of genes involved in inflammation and apoptosis.

**Figure 8.**
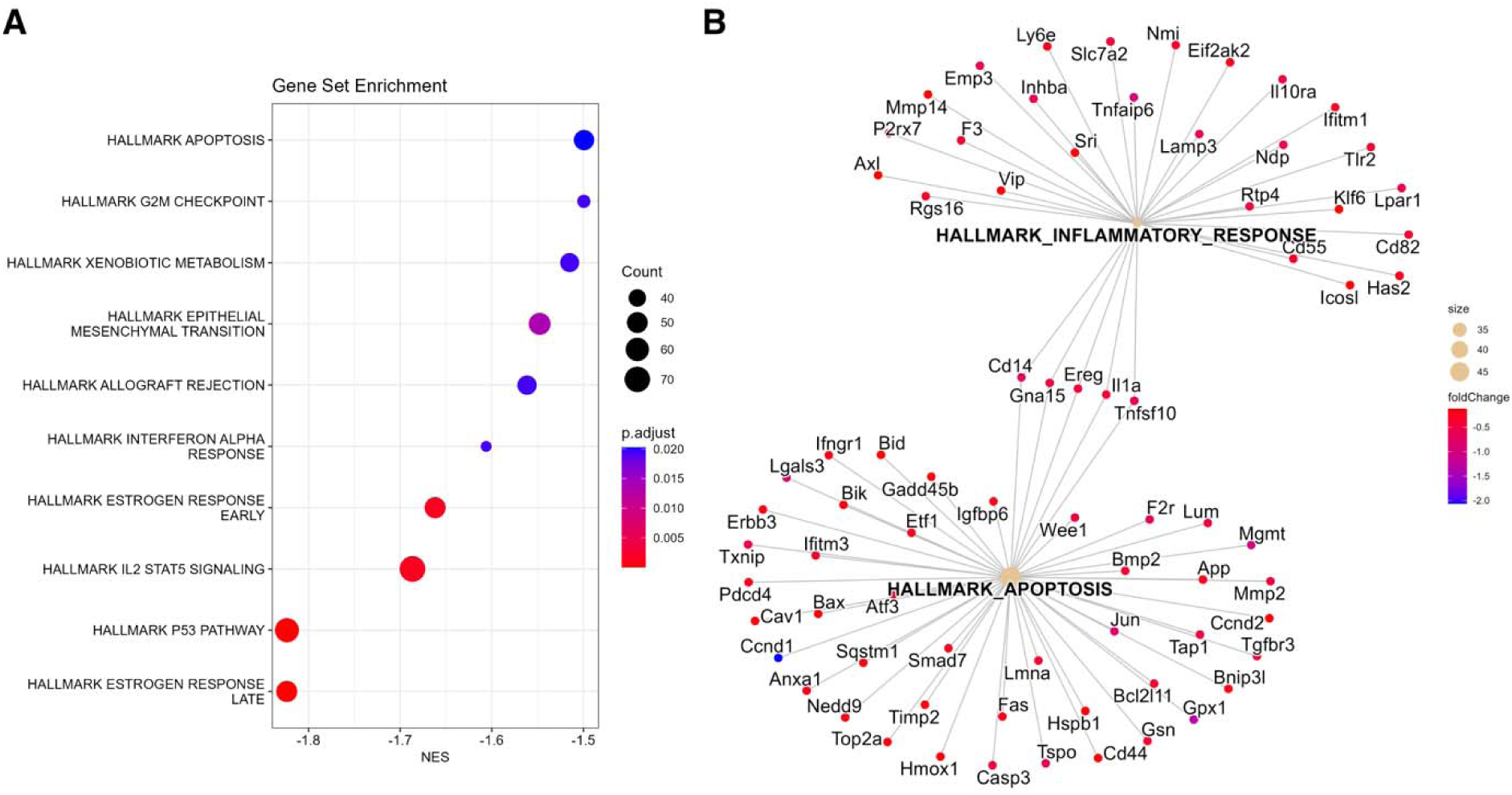
Transcriptomic analysis of diabetic mice given intranasal insulin vs intranasal saline. A. Pathway enrichment analysis using hallmark gene set with gene set enrichment analysis (GSEA) revealed ten significantly downregulated pathways in the intranasal insulin group. B. Network map of enriched pathways of inflammation and apoptosis further elucidated the differentially regulated genes.

## Discussion

To the best of our knowledge, we have demonstrated for the first time the successful delivery of insulin to the retina via the intranasal route. A PubMed search dated Jan 1, 1970 to Jun 30, 2025 with the terms “intranasal insulin” and “retina” yielded 0 results. Retinal insulin immunostaining in the group treated with insulin was much higher than that for both the wildtype and diabetic groups treated with saline. Our results also showed that insulin delivered intranasally successfully prevented reductions in b-wave and oscillatory potential amplitudes in diabetic mice, potentially by decreased glial activation, decreased cell death and downregulation of multiple inflammatory and apoptotic pathways. We found no reduction of blood glucose levels after administration of intranasal insulin and thus, further verified that intranasal insulin exerted its effects centrally and not peripherally since it did not lower serum glycemic levels.

Neurodegenerative changes have been observed in the retina of diabetic patients in the absence of any diabetic retinopathy.^21^ Moreover, observational reports have shown that neurodegeneration measured by multifocal ERG can predict which locations would develop DR in the future.^22–24^ Specifically, diabetic patients without retinopathy first present with reductions in their ERG b-wave response.^25^ This decline in inner retinal function is a hallmark of early disease and recapitulated in animal models of diabetes, including ours. In our study, *db/db* mice treated with saline had significant reductions in b-wave and oscillatory potential amplitudes. These studies are consistent with those shown in other studies using *db/db* mice,^26^ Ins2Akita mice^27^ and streptozocin induced rodent models.^28^ In light of these findings, our results showing that insulin, given intranasally every day for 10 weeks, can prevent reductions in b-wave and oscillatory potential amplitudes in diabetes are highly novel.

The ERG b-wave is thought to arise from bipolar cells, Muller glia or both,^29^ while oscillatory potentials are thought to largely be generated by interactions among inner retina neurons, in particular amacrine cells.^30^ Structurally, these changes in diabetes have corresponded to inner retinal thinning involving the ganglion cell layer, retinal nerve fiber layer, inner plexiform layer and inner nuclear layer in both human and animal studies.^28,31,32^ In our study, inner retinal thickness was also decreased in the *db/db* saline group. Treatment with insulin did not appear to reduce this loss of inner retinal thickness in the diabetic group. However, our study revealed a decrease in outer retinal thickness in the *db/db* saline group compared to control mice, and that treatment with insulin did prevent this reduction. Reductions in outer retinal thickness in animal models of DR^28^ have been shown previously and correlated with more advanced stages of diabetic retinopathy in humans.^33^ Since inner retinal thinning precedes outer retinal thinning in diabetes, we believe our results suggest that intranasal insulin over 10 weeks may not have sufficiently prevented inner retinal thinning which occurs earlier in diabetes, but was able to reduce outer retinal thinning which occurs later in the disease.

Reactive gliosis and apoptosis are two hallmark processes of neurodegeneration. As such, we evaluated GFAP, expressed in astrocytes and reactive Muller cells in response to injury and inflammation. Consistent with other reports,^26,34^ our results showed that diabetes upregulated GFAP expression in the retina. Our results additionally demonstrated that the diabetic group treated with insulin had less GFAP expression compared to the wildtype control and diabetic mice treated with saline. We also quantified cleaved caspase 3 and TUNEL immunoreactivity to examine apoptosis. When compared to the wildtype controls, higher number of cells with caspase 3 immunoreactivity were counted in diabetic mice treated with saline but not in diabetic mice treated with insulin. There was also a near significant trend of a higher count of TUNEL positive cells in the diabetic group treated with saline which was not seen in the diabetic group treated with insulin. Our results suggest that intranasal insulin treatment reduces glial activation and apoptosis in the retina of diabetic mice.

When we performed transcriptomic analysis of the retina, we found that insulin reduced expression of several genes involved in inflammation and apoptosis in the diabetic retina. For example, Bax (Bcl-2-Associated X protein) is a pro-apoptotic cytokine that has been found to be elevated in the postmortem retinas of diabetic donors when compared to controls.^35^ Bax was localized to vascular and neural cells of the inner retina, and was highly expressed in retinal vascular pericytes and outer cells of retinal capillaries in conjunction with DNA fragmentation in diabetes.^35^ Caspase 3 is another pro-apoptotic enzyme that has been shown to accelerate cell death of endothelial cells and pericytes in DR. Additionally, HMOX-1 (heme-oxygenase 1) is a ferroptosis related gene that was found to be differentially expressed in DR.^36^ Ferroptosis is programmed cell death that encompasses high iron-dependent lipid peroxidation, and has been implicated in DR initiation and progression. Given the role of Bax, caspase 3 and HMOX-1 in mediating accelerated apoptosis of retinal neural and vascular cells in diabetes, our finding that intranasal insulin can downregulate their expression is significant and represents a mechanistic pathway through which intranasal insulin can exert its neuroprotective effects. Meanwhile, IL-1alpha has been shown to stimulate angiogenesis by activating the VEGF-VEGFR2 signaling pathway.^37^ Our findings of downregulation of this and other apoptotic and pro-inflammatory genes demonstrates the potential efficacy of intranasal insulin in modifying signaling pathways through differential gene expression.

Taken together, these results suggest that intranasal insulin, a non-invasive treatment, exerted neuroprotective effects in the retina of diabetic *db/db* mice after once daily treatments over 10 weeks by modifying differential expression of inflammatory and apoptotic genes. Importantly, intranasal insulin did not lower peripheral blood glucose levels and exerted its neuroprotective effects directly on the retina. Insulin signaling occurs through its transmembrane receptors, which have been found to be widely expressed in the human retina, most abundantly in neuronal cell bodies, including photoreceptors, and in the plexiform layer and Muller cells.^38^

Recent evidence has also shown that insulin is produced locally in the neuroretina and RPE. mRNA transcripts of insulin 1 (Ins1) and insulin 2 (Ins2) were detected in the ganglion cell layer, inner nuclear layer, outer nuclear layer and retinal pigment epithelium in murine retinas.^39^ Similarly, retinal transcripts of insulin were seen in the ganglion cell layer, inner nuclear layer, inner segment of photoreceptors and retinal pigment epithelium in human donors.^39^ These findings suggest that the high basal insulin receptor signaling in the retina is driven by local insulin production in addition to contributions from peripheral circulating insulin.

Diabetes has been shown to disrupt insulin receptor signaling and local insulin production in the retina. Insulin receptors autophosphorylate and activate downstream signaling kinases.^5^ After 4 weeks of streptozocin induced diabetes, constitutive insulin receptor kinase activity was reduced and after 12 weeks, further loss of insulin receptor autophosphorylation, expression and activity was observed.^5^ Delivery of both systemic and intravitreal exogenous insulin restored these deficits in insulin receptor signaling.^5^ Meanwhile, local insulin production has been shown to be upregulated in response to acute stress but downregulated in chronic disease like diabetes. After 8 weeks of streptozocin induced diabetes, Ins1 and Ins2 expression was increased. However, after 23 weeks of diabetes, Ins1 expression was downregulated while Ins2 expression remained comparable to control mice. In human diabetic donors, there was a marked decrease in insulin transcripts in the RPE, but not the neuroretina. Both retinal neurons and vascular endothelial cells have been found to be dependent on the insulin mediated phosphatidylinositol 3-kinase/Akt mechanism for survival, and deficiency in local insulin production and disruption of insulin receptor signaling in diabetes is thought to induce apoptosis in both these cell types.^6,40^

Insulin’s role in preventing neuronal cell death in the retina has been well described. In differentiated R28 cells, which model retinal neurons, treatment with insulin reduces apoptosis by activating the phosphatidylinositol 3-kinase/Akt pathway and reducing caspase 3 activation.^6^ In streptozocin-diabetic rats, insulin delivery through a subcutaneous implant (2-4U daily) for 1 month successfully reduced the number of TUNEL-HRP positive cells in retinal wholemounts.^41^ The same group also demonstrated that 48 hours of subcutaneous 10U twice daily insulin treatment improved tight junction protein expression and glial reactivity in the retina.^42^ More recently, Rong and coauthors performed subconjunctival injections of insulin-loaded chitosan nanoparticles entrapped in a hydrogel in streptozocin-diabetic rats and showed a decrease in the reduction of b-waves, retinal microstructural changes and retinal apoptosis after two weeks of therapy.^43^

In contrast, in another study using streptozocin-diabetic rats, insulin delivery via a subcutaneous implant (2U insulin daily over 40 days) exacerbated blood retinal barrier breakdown via elevated VEGF and hypoxia inducible factor (HIF) α expression.^44^ These results were hypothesized to explain acute worsening of diabetic retinopathy in patients who commence high dose insulin. However, published literature on acute DR worsening after institution of insulin treatment in humans remains controversial. In a meta-analysis of seven cohort studies, the authors concluded that the association between insulin use and DR in patients with type 2 diabetes became non-significant when the data was adjusted for duration of diabetes.^45^ Still, upregulation of VEGF production after insulin treatment has been shown in several cell types including vascular smooth muscle cells,^46^ cardiomyocytes^47^ and fibroblasts^48^.

Insulin has pleiotropic effects and may respond differently to various stages of stress. We postulate that acute stress causes increased production of endogenous insulin, and when combined with exogenous insulin, the high levels of insulin can increase VEGF and HIF α production. Increased vascularization can be beneficial in certain contexts, such as wound healing enhancement, transplant acceptance and revascularization after myocardial ischemia.^49^ However, in other settings, this can be pathologic, for example in the retina, where short term worsening of diabetic retinopathy has been observed acutely after insulin therapy. However, in chronic stress, where there is deficient local insulin production, exogenous insulin’s neurotrophic effects may predominate, resulting in less progression of diabetic retinopathy. For example, in DCCT, patients with type 1 diabetes (n=1441, average 6.5 years of follow up) given intensive insulin therapy had reduced development of new diabetic retinopathy (for those with no retinopathy), and less progression to worse diabetic retinopathy (for those with baseline mild retinopathy) compared to patients given conventional insulin therapy.^50^ In the ACCORD Eye Study, patients with type 2 diabetes (n=2856 with 4 year data) given intensive insulin therapy similarly had less progression of diabetic retinopathy.^51^

Currently, available methods of delivering insulin to the retina include systemic, intravitreal and subconjunctival approaches, which as previously discussed, has been shown to decrease neurodegeneration associated with diabetes.^6,8,9^ However, there are numerous challenges associated with current routes of insulin delivery. Insulin has a short half-life in the plasma,^12^ and high levels of systemic insulin would have to be administered to reduce the risk of retinopathy, putting the patients at risk for hypoglycemia.^13^ Due to corneal and conjunctival barriers, and rapid precorneal tear loss, topical eye drops are an inefficient way to treat retinal diseases.^14,15^ Intravitreal and subconjunctival injections are invasive, requires frequent visits to the treating ophthalmologist creating substantial treatment burden to the healthcare system, and risk of endophthalmitis although low, is not negligible.^9,16^ In contrast, intranasal insulin administration, as used in our study, is non-invasive, can be easily self-administered, avoids hepatic first-pass elimination and reaches the retina and the brain without impacting peripheral circulating insulin levels or causing hypoglycemia.^17,18^

Our results show, for the first time, that insulin delivered intranasally reaches the retina and has neuroprotective properties in diabetic murine retina. These results suggest the feasibility of utilizing a novel method of insulin delivery to the retina that offers many advantages over current routes of delivery. These results also offer proof of concept for the potential intranasal delivery of other therapeutics to the retina, which may ultimately be transformative to the field of ophthalmology and medicine.

## Acknowledgements

The authors gratefully thank Ryan G. Phillips, Jonathan Groothoff, Sean Wang, Rameen Janjua, Ava Peterson and Tian Yuan for their generous technical support.

